# Transmissibility, infectivity, and immune resistance of the SARS-CoV-2 BA.2.86 variant

**DOI:** 10.1101/2023.09.07.556636

**Authors:** Keiya Uriu, Jumpei Ito, Yusuke Kosugi, Yuri L Tanaka, Yuka Mugita, Ziyi Guo, Alfredo A Hinay, Olivia Putri, Yoonjin Kim, Ryo Shimizu, MST Monira Begum, Michael Jonathan, The Genotype to Phenotype Japan (G2P-Japan) Consortium, Akatsuki Saito, Terumasa Ikeda, Kei Sato

## Abstract

In September 2023, the SARS-CoV-2 XBB descendants, such as XBB.1.5 and EG.5.1 (originally XBB.1.9.2.5.1), are predominantly circulating worldwide. Unexpectedly, however, a lineage distinct from XBB was identified and named BA.2.86 on August 14, 2023. Notably, BA.2.86 bears more than 30 mutations in the spike (S) protein when compared to XBB and the parental BA.2, and many of them are assumed to be associated with immune evasion. Although the number of reported cases is low (68 sequences have been reported as of 7 September 2023), BA.2.86 has been detected in several continents (Europe, North America and Africa), suggesting that this variant may be spreading silently worldwide. On 17 August 2023, the WHO designated BA.2.86 as a variant under monitoring. Here we show evidence suggesting that BA.2.86 potentially has greater fitness than current circulating XBB variants including EG.5.1. The pseudovirus assay showed that the infectivity of BA.2.86 was significantly lower than that of B.1.1 and EG.5.1, suggesting that the increased fitness of BA.2.86 is not due to the increased infectivity. We then performed a neutralization assay using XBB breakthrough infection sera to address whether BA.2.86 evades the antiviral effect of the humoral immunity induced by XBB subvariants. The 50% neutralization titer of XBB BTI sera against BA.2.86 was significantly (1.6-fold) lower than those against EG.5.1. The sera obtained from individuals vaccinated with 3rd-dose monovalent, 4th-dose monovalent, BA.1 bivalent, and BA.5 bivalent mRNA vaccines exhibited very little or no antiviral effects against BA.2.86. Moreover, the three monoclonal antibodies (Bebtelovimab, Sotrovimab and Cilgavimab), which worked against the parental BA.2, did not exhibit antiviral effects against BA.2.86. These results suggest that BA.2.86 is one of the most highly immune evasive variants ever.

## Text

In September 2023, the SARS-CoV-2 XBB descendants, such as XBB.1.5 and EG.5.1 (originally XBB.1.9.2.5.1), are predominantly circulating worldwide according to Nextstrain (https://nextstrain.org/ncov/gisaid/global/6m). Unexpectedly, however, a lineage distinct from XBB was identified and named BA.2.86 on August 14, 2023.^1^ Notably, BA.2.86 bears more than 30 mutations in the spike (S) protein when compared to XBB and the parental BA.2, and many of them are assumed to be associated with immune evasion (**Figure 1A**).^1^ Although the number of reported cases is low (68 sequences have been reported as of 7 September 2023), BA.2.86 has been detected in several continents (Europe, North America and Africa), suggesting that this variant may be spreading silently worldwide. On 17 August 2023, the WHO designated BA.2.86 as a variant under monitoring.^2^

**Figure 1.**
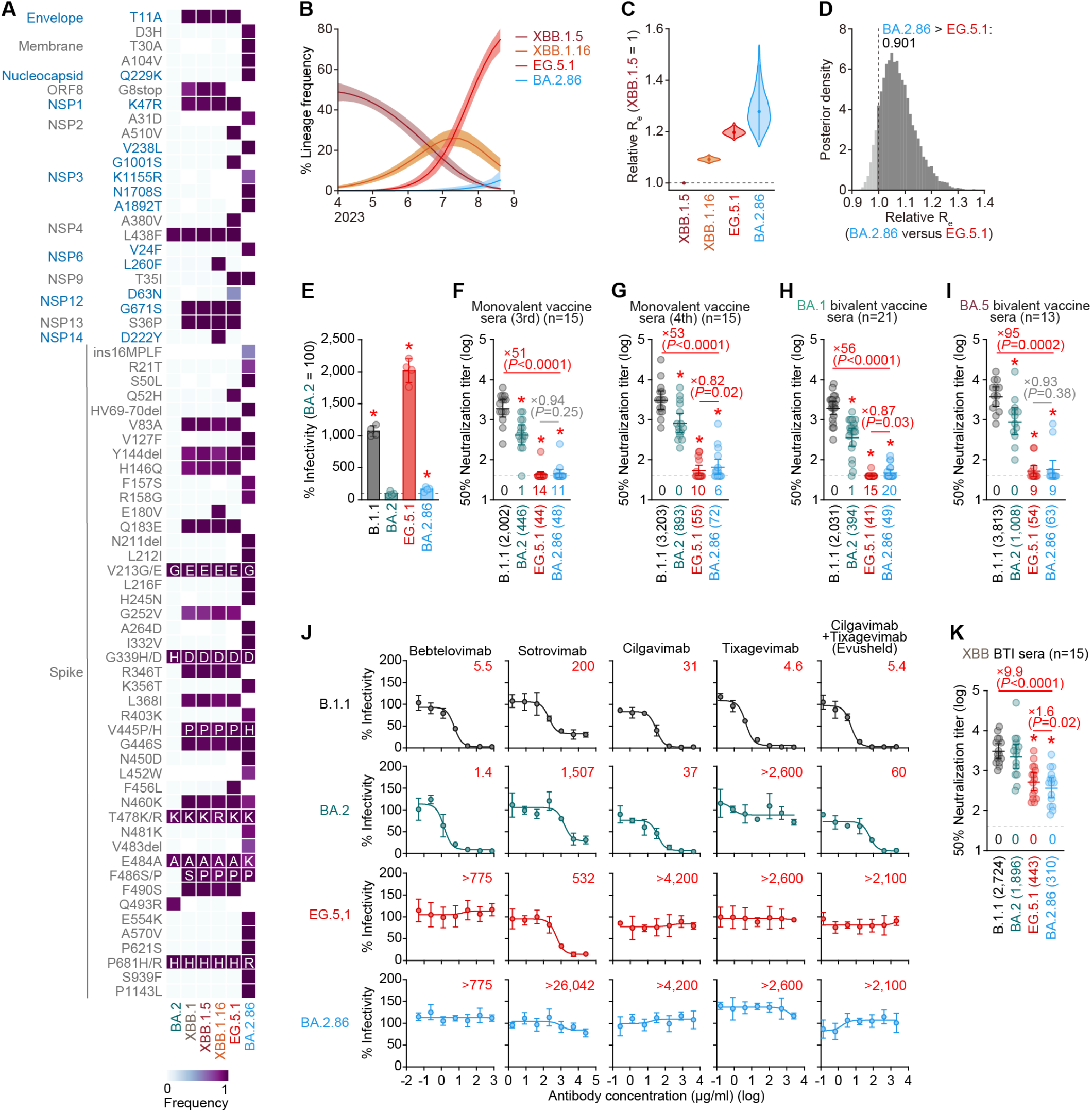
Virological features of BA.2.86. (**A**) Frequency of mutations of interest in BA.2.86 and the representative XBB sublineages. Only mutations with a frequency >0.5 in at least one but not all the lineages are shown. (**B**) Estimated epidemic dynamics of BA.2.86 and the representative XBB sublineages in Denmark from April 1, 2023 to September 4, 2023. Line, posterior mean; ribbon, 95% Bayesian confidence interval (CI). (**C**) Estimated relative R_e_ of BA.2.86 and the representative XBB sublineages in Denmark. The relative R_e_ of XBB.1.5 is set to 1 (horizontal dashed line). Violin represents the posterior distribution encompassed by the 99% Bayesian CI. The dot denotes the posterior mean, and the line highlights the 95% Bayesian CI. (**D**) The posterior distribution of the R_e_ of BA.2.86 relative to that of EG.5.1. Posterior probability that the R_e_ of BA.2.86 exceeds that of EG.5.1 is denoted. (**E**) Lentivirus-based pseudovirus assay. HOS-ACE2/TMPRSS2 cells were infected with pseudoviruses bearing each S protein of B.1.1, BA.2, EG.5.1, and BA.2.86. The amount of input virus was normalized to the amount of HIV-1 p24 capsid protein. The percentage infectivity of B.1.1, EG.5.1, and BA.2.86 compared to that of BA.2 are shown. The horizontal dash line indicates the mean value of the percentage infectivity of the BA.2. Assays were performed in quadruplicate, and a representative result of six independent assays is shown. The presented data are expressed as the average ± standard deviation. Each dot indicates the result of an individual replicate. Statistically significant differences (*, p < 0.05) versus BA.2 were determined by two-sided Student’s *t* tests. Red asterisks indicate an increased percentage of infectivity. (**F-I**) Neutralization assay with vaccine sera. Assays were performed with pseudoviruses harboring the S proteins of B.1.1, BA.2, EG.5.1, or BA.2.86. In (**F**) to (**I**), the following sera were used: 3rd-dose monovalent vaccine sera (15 donors) (**F**), 4th-dose monovalent vaccine sera (15 donors) (**G**), BA.1 bivalent vaccine (21 donors) (**H**), and BA.5 bivalent vaccine (13 donors) (**I**). Vaccine sera were collected at 1 month after the last vaccination. (**J**) Neutralization assay with monoclonal antibodies. Four therapeutic monoclonal antibodies (bebtelovimab, sotrovimab, cilgavimab, and tixagevimab) and one combination, cilgavimab+tixagevimab (Evusheld), were tested. The assay of each antibody was performed in triplicate at each concentration to determine the 50% neutralization concentration (μg/ml). The presented data are expressed as the average with standard deviation, and representative neutralization curves are shown. The red numbers in the panels indicate the 50% neutralization concentration (μg/ml). (**K**) Neutralization assay with convalescent sera. Convalescent sera from fully vaccinated individuals who had been infected with XBB.1.5 (four 3-dose vaccinated donors and two 5-dose vaccinated donors. 6 donors in total), XBB.1.9 (one 2-dose vaccinated donor, one 4-dose vaccinated donors, and two 5-dose vaccinated donors. 4 donors in total), and XBB.1.16 (three 3-dose vaccinated donors, one 4-dose vaccinated donors, and one 6-dose vaccinated donor. 5 donors in total). In (**F**-**I**) and (**K**), assays for each serum sample were performed in triplicate to determine the 50% neutralization titer (NT_50_). Each dot represents one NT_50_ value, and the geometric mean and 95% confidence interval are shown. The number in parenthesis indicates the geometric mean of NT_50_ values. The horizontal dashed line indicates the detection limit (40-fold). The number of serum donors with the 50% NT_50_ values below the detection limit (40-fold dilution) is shown in the figure (under the bars and dots of each variant). Statistically significant differences (*, P < 0.05) versus B.1.1 were determined by two-sided Wilcoxon signed-rank tests and indicated with asterisks. Red asterisks indicate decreased NT_50_ values. The fold changes of reciprocal NT_50_ are calculated between B.1.1 and BA.2.86 and that between EG.5.1 and BA.2.86. Background information on the convalescent donors is summarized in **Table S1**.

We first estimated the effective reproduction number (R_e_) of BA.2.86 based on the genome surveillance data until September 4, 2023, in Denmark, where multiple XBB subvariants including EG.5.1 are currently co-circulating, and twelve BA.2.86 sequences were reported (**Figure 1B**). Although there was considerable uncertainty in this estimation due to a smaller number of available BA.2.86 sequences, the R_e_ of BA.2.86 was 1.29-fold greater than that of XBB.1.5 (Bayesian confidence interval: 1.17–1.47) (**Figure 1C**). Importantly, the R_e_ of BA.2.86 was comparable to or even greater than that of EG.5.1, one of the most rapidly expanding XBB subvariants (**Figure 1C**), and the estimated posterior probability that the R_e_ of BA.2.86 exceeds that of EG.5.1 was 0.901 (**Figure 1D**). These findings suggest that BA.2.86 potentially has greater fitness than current circulating XBB variants including EG.5.1.

To assess the possibility that the enhanced infectivity of BA.2.86 contributes to its augmented R_e_, we prepared the lentivirus-based pseudoviruses with the S proteins of BA.2.86, EG.5.1, parental BA.2, and ancestral D614G-bearing B.1.1. The pseudovirus assay showed that the infectivity of BA.2.86 was significantly lower than that of B.1.1 and EG.5.1 (**Figure 1E**), suggesting that the increased R_e_ of BA.2.86 is not due to the increased infectivity.

We then performed neutralization assays using vaccine sera to assess the possibility that BA.2.86 evades the antiviral effect of vaccine-induced humoral immunity. The sera obtained from individuals vaccinated with 3rd-dose monovalent, 4th-dose monovalent, BA.1 bivalent, and BA.5 bivalent mRNA vaccines exhibited very little or no antiviral effects against BA.2.86 (**Figures 1G-1J**). Additionally, the three monoclonal antibodies (Bebtelovimab, Sotrovimab and Cilgavimab), which worked against the parental BA.2,^3^ did not exhibit antiviral effects against BA.2.86 (**Figure 1J**). Finally, a neutralization assay using XBB breakthrough infection sera showed that the 50% neutralization titer of XBB breakthrough infection sera against BA.2.86 was significantly (1.6-fold) lower than those against EG.5.1 (p<0.0001) (**Figure 1K**). Altogether, these results suggest that BA.2.86 is one of the most highly immune evasive variants ever.

## Grants

Supported in part by AMED SCARDA Japan Initiative for World-leading Vaccine Research and Development Centers “UTOPIA” (JP223fa627001, to Kei Sato), AMED SCARDA Program on R&D of new generation vaccine including new modality application (JP223fa727002, to Kei Sato); AMED Research Program on Emerging and Re-emerging Infectious Diseases (JP22fk0108146, to Kei Sato; JP21fk0108494 to G2P-Japan Consortium, Terumasa Ikeda and Kei Sato; JP21fk0108425, to Kei Sato; JP21fk0108432, to Kei Sato; JP22fk0108511, to G2P-Japan Consortium, Akatsuki Saito, Terumasa Ikeda and Kei Sato; JP22fk0108516, to Kei Sato; JP22fk0108506, to Akatsuki Saito and Kei Sato; JP22fk0108534, to Terumasa Ikeda and Kei Sato); AMED Research Program on HIV/AIDS (JP23fk0410047, to Akatsuki Saito; JP 23fk0410056, to Akatsuki Saito; JP23fk0410058, to Akatsuki Saito; JP22fk0410055, to Terumasa Ikeda; JP22fk0410039, to Kei Sato); JST PRESTO (JPMJPR22R1, to Jumpei Ito); JST CREST (JPMJCR20H4, to Kei Sato); JSPS KAKENHI Grant-in-Aid for Scientific Research C (22K07103, to Terumasa Ikeda); JSPS KAKENHI Grant-in-Aid for Early-Career Scientists (23K14526, to Jumpei Ito); JSPS Leading Initiative for Excellent Young Researchers (LEADER) (to Terumasa Ikeda); JSPS Core-to-Core Program (A. Advanced Research Networks) (JPJSCCA20190008, Kei Sato); JSPS Research Fellow DC2 (22J11578, to Keiya Uriu); JSPS Research Fellow DC1 (23KJ0710, to Yusuke Kosugi); The Tokyo Biochemical Research Foundation (to Kei Sato); The Mitsubishi Foundation (to Kei Sato); Mochida Memorial Foundation for Medical and Pharmaceutical Research (to Terumasa Ikeda); The Naito Foundation (to Terumasa Ikeda) ; International Joint Research Project of the Institute of Medical Science, the University of Tokyo (to Terumasa Ikeda and Akatsuki Saito).

## Declaration of interest

J.I. has consulting fees and honoraria for lectures from Takeda Pharmaceutical Co. Ltd. K.S. has consulting fees from Moderna Japan Co., Ltd. and Takeda Pharmaceutical Co. Ltd. and honoraria for lectures from Gilead Sciences, Inc., Moderna Japan Co., Ltd., and Shionogi & Co., Ltd. The other authors declare no competing interests. All authors have submitted the ICMJE Form for Disclosure of Potential Conflicts of Interest. Conflicts that the editors consider relevant to the content of the manuscript have been disclosed.

## Materials and Methods

### Ethics statement

All protocols involving specimens from human subjects recruited at Kyoto University and Interpark Kuramochi Clinic were reviewed and approved by the Institutional Review Boards of Kyoto University (approval ID: G1309) and Interpark Kuramochi Clinic (approval ID: G2021-004). All human subjects provided written informed consent. All protocols for the use of human specimens were reviewed and approved by the Institutional Review Boards of The Institute of Medical Science, The University of Tokyo (approval IDs: 2021-1-0416 and 2021-18-0617), Kumamoto University (approval ID: 2066), and University of Miyazaki (approval ID: O-1021).

### Human serum collection

Convalescent sera were collected from fully vaccinated individuals who had been infected with XBB sublineages [XBB.1.5 (four 3-dose vaccinated, two 5-dose vaccinated; time interval between the last vaccination and infection, 114-435 days; 14-46 days after testing. n=6 in total; average age: 42 years, range: 15-74 years, 16.7% male), XBB.1.9 (one 2-dose vaccinated, one 4-dose vaccinated and two 5-dose vaccinated; time interval between the last vaccination and infection, 176–248 days; 13–32 days after testing. n=4 in total; average age: 56 years, range: 21–88 years, 25% male), and XBB.1.16 (three 3-dose vaccinated, one 4-dose vaccinated and one 6-dose vaccinated; time interval between the last vaccination and infection, 35–481 days; 8–30 days after testing. n=5 in total; average age: 41 years, range: 26–56 years, 80% male)]. Vaccine sera of fifteen individuals who had BNT162b2 vaccine (Pfizer/BioNTech) (average age: 38 years, range: 24–48 years; 53% male) were obtained at one month after the third dose. Vaccine sera from fifteen individuals who had been vaccinated with BNT162b2 vaccine (Pfizer/BioNTech) (average age: 42 years, range: 30–56 years, 40% male) were obtained at one month after the fourth dose. Vaccine sera from twenty-one individuals who had been vaccinated with BA.1 bivalent vaccine (average age: 55 years, range: 30–80 years, 38% male) were obtained at one month after the fourth dose. Vaccine sera of thirteen individuals who had BA.5 bivalent vaccine (average age: 55 years, range: 27–86 years, 54% male) were obtained at one month after the last vaccination. The SARS-CoV-2 variants were identified as previously described.^1-3^ Sera were inactivated at 56°C for 30 minutes and stored at –80°C until use. The details of the convalescent sera are summarized in **Table S1**.

### Epidemic dynamics analysis and mutation frequency calculation

In the present study, we analyzed the viral genomic surveillance data deposited in the GISAID database (https://www.gisaid.org/; release date: September 4, 2023). We used the data from April 1, 2023 in this analysis. We excluded the sequence records with the following features: i) a lack of collection date information; ii) sampling in animals other than humans; iii) sampling by quarantine; and iv) without the PANGO lineage information. We modeled the epidemic dynamics of viral lineages in Denmark, where 12 sequences of BA.2.86 were detected by September 4, 2023. Importantly, this country did not employ biased sequencing strategy for BA.2.86. In the downstream analysis, we only used sequences for BA.2.86 or PANGO lineages with >50 sequences in the dataset. We counted the daily frequency of each viral lineage. Subsequently, epidemic dynamics and relative R_e_ value for each viral lineage were estimated according to the Bayesian multinomial logistic model, described in our previous study.^2^ Briefly, we estimated the logistic slope parameter β_*l*_ for each viral lineage using the model and then calculated relative R_e_ for each lineage *r*_*l*_ as *r*_*l*_ = *exp*(γβ_*l*_) where γ is the average viral generation time (2.1 days) (http://sonorouschocolate.com/covid19/index.php?title=Estimating_Generation_Time_Of_Omicron). For parameter estimation, the intercept and slope parameters of XBB.1.5 were fixed at 0. Consequently, the relative R_e_ of XBB.1.5 was fixed at 1, and those of the other lineages were estimated relative to that of XBB.1.5. Parameter estimation was performed via the MCMC approach implemented in CmdStan v2.30.1 (https://mc-stan.org) with CmdStanr v0.5.3 (https://mc-stan.org/cmdstanr/). Four independent MCMC chains were run with 1,000 and 4,000 steps in the warmup and sampling iterations, respectively. We confirmed that all estimated parameters showed <1.01 R-hat convergence diagnostic values and >200 effective sampling size values, indicating that the MCMC runs were successfully convergent. Information on the estimated parameters is summarized in **Table S2**. In **Figures 1B and 1C**, results for XBB.1.5, XBB.1.16, EG.5.1, and BA.2.86 are shown.

### Cell culture

The Lenti-X 293T cell line (Takara, Cat# 632180), HEK293T cells (a human embryonic kidney cell line; ATCC, CRL-3216) and HOS-ACE2/TMPRSS2 cells (kindly provided by Dr. Kenzo Tokunaga), a derivative of HOS cells (a human osteosarcoma cell line; ATCC CRL-1543) stably expressing human ACE2 and TMPRSS2,^12,13^ were maintained in Dulbecco’s modified Eagle’s medium (DMEM) (high glucose) (Wako, Cat# 044-29765) containing 10% fetal bovine serum (Sigma-Aldrich Cat# 172012-500ML), 100 units penicillin and 100 ug/ml streptomycin (Sigma-Aldrich, Cat# P4333-100ML).

### Plasmid construction

Plasmids expressing the SARS-CoV-2 spike (S) proteins of the ancestral B.1.1 (D614G-bearing virus), Omicron BA.2, and EG.5.1 were prepared in our previous studies.^1,2,4-9^ Plasmids expressing the S protein of a BA.2.86 sequence (GISAID ID: EPI_ISL_18114953) was generated by site-directed overlap extension PCR using pC-SARS2-S BA.2^1^ as the template and the primers listed in **Table S3**. The resulting PCR fragment was subcloned into the KpnI-NotI site of the pCAGGS vector^10^ using In-Fusion HD Cloning Kit (Takara, Cat# Z9650N). Nucleotide sequences were determined by DNA sequencing services (Eurofins), and the sequence data were analyzed by SnapGene software v6.1.1 (www.snapgene.com).

### Preparation of monoclonal antibodies

Four monoclonal antibodies (bebtelovimab, sotrovimab, cilgavimab, tixagevimab) were prepared as previously described.^3,11^ Briefly, the pCAGGS vectors containing the sequences encoding the immunoglobulin heavy and light chains were cotransfected into HEK293T cells at 1:1 ratio using PEI Max (Polysciences, Cat# 24765-1). The culture medium was refreshed with DMEM (low glucose) (Wako, Cat# 041-29775) containing 10% FBS without PS. At 96 h posttransfection, the culture medium was harvested, and the antibodies were purified using NAb protein A plus spin kit (Thermo Fisher Scientific, Cat# 89948) according to the manufacturer’s protocol.

### Pseudovirus preparation

Pseudoviruses were prepared as previously described.^8^ Briefly, lentivirus (HIV-1)-based, luciferase-expressing reporter viruses were pseudotyped with the SARS-CoV-2 S protein. One prior day of transfection, the LentiX-293T or HEK293T cells were seeded at a density of 2 × 10^6^ cells. The LentiX-293T or HEK293T cells were cotransfected with 1 μg psPAX2-IN/HiBiT (a packaging plasmid encoding the HiBiT-tag-fused integrase^14^), 1 μg pWPI-Luc2 (a reporter plasmid encoding a firefly luciferase gene^14^) and 500 ng plasmids expressing parental S protein or its derivatives using TransIT-293 transfection reagent (Mirus, Cat# MIR2704) or TransIT-LT1 (Takara, Cat# MIR2300) according to the manufacturer’s protocol. Two days post transfection, the culture supernatants were harvested and filtrated. The amount of produced pseudovirus particles was quantified by the HiBiT assay using Nano Glo HiBiT lytic detection system (Promega, Cat# N3040) as previously described^14^. In this system, HiBiT peptide is produced with HIV-1 integrase and forms NanoLuc luciferase with LgBiT, which is supplemented with substrates. In each pseudovirus particle, the detected HiBiT value is correlated with the amount of the pseudovirus capsid protein, HIV-1 p24 protein.^14^ Therefore, we calculated the amount of HIV-1 p24 capsid protein based on the HiBiT value measured, according to the previous paper.^14^ To measure viral infectivity, the same amount of pseudovirus normalized with the HIV-1 p24 capsid protein was inoculated into HOS-ACE2/TMPRSS2 cells. At two days postinfection, the infected cells were lysed with a Bright-Glo luciferase assay system (Promega, Cat# E2620), and the luminescent signal produced by firefly luciferase reaction was measured using a GloMax explorer multimode microplate reader 3500 (Promega) or CentroXS3 LB960 (Berthhold Technologies). The pseudoviruses were stored at –80°C until use.

### Neutralization assay

Neutralization assays were performed as previously described.^8^ The SARS-CoV-2 S pseudoviruses (counting ∼50,000 relative light units) were incubated with serially diluted (40-fold to 29,160-fold dilution at the final concentration) heat-inactivated sera at 37°C for 1 hour. Pseudoviruses without sera were included as controls. Then, 20 μl mixture of pseudovirus and serum was added to HOS-ACE2/TMPRSS2 cells (10,000 cells/100 μl) in a 96-well white plate. Two days post infection, the infected cells were lysed with a Bright-Glo luciferase assay system (Promega, Cat# E2620), and the luminescent signal was measured using a GloMax explorer multimode microplate reader 3500 (Promega). The assay of each serum sample was performed in triplicate, and the 50% neutralization titer (NT_50_) was calculated using Prism 9 (GraphPad Software).

## Data availability

Dataset used in the epidemic dynamics analysis in this study is available from the GISAID database (https://www.gisaid.org; EPI_SET_230907ur). The GISAID supplemental tables for EPI_SET_230907ur is available in the GitHub repository (https://github.com/TheSatoLab/BA.2.86_short).

## Consortia

### The Genotype to Phenotype Japan (G2P-Japan) Consortium

**The Institute of Medical Science, The University of Tokyo, Japan**

Yu Kaku, Naoko Misawa, Arnon Plianchaisuk, Jarel Elgin M. Tolentino, Luo Chen, Shigeru Fujita, Lin Pan, Mai Suganami, Mika Chiba, Ryo Yoshimura, Kyoko Yasuda, Keiko Iida, Naomi Ohsumi, Adam P. Strange, Shiho Tanaka, Kaho Okumura

**Hokkaido University, Japan**

Takasuke Fukuhara, Tomokazu Tamura, Rigel Suzuki, Saori Suzuki, Hayato Ito, Keita Matsuno, Hirofumi Sawa, Naganori Nao, Shinya Tanaka, Masumi Tsuda, Lei Wang, Yoshikata Oda, Zannatul Ferdous, Kenji Shishido

**Tokai University, Japan**

So Nakagawa

**Kyoto University, Japan**

Kotaro Shirakawa, Akifumi Takaori-Kondo, Kayoko Nagata, Ryosuke Nomura, Yoshihito Horisawa, Yusuke Tashiro, Yugo Kawai, Kazuo Takayama, Rina Hashimoto, Sayaka Deguchi, Yukio Watanabe, Ayaka Sakamoto, Naoko Yasuhara, Takao Hashiguchi, Tateki Suzuki, Kanako Kimura, Jiei Sasaki, Yukari Nakajima, Hisano Yajima

**Hiroshima University, Japan**

Takashi Irie, Ryoko Kawabata

**Kyushu University, Japan**

Kaori Tabata

**Kumamoto University, Japan**

Hesham Nasser, Otowa Takahashi, Kimiko Ichihara, Takamasa Ueno, Chihiro Motozono, Mako Toyoda

**University of Miyazaki, Japan**

Maya Shofa, Yuki Shibatani, Tomoko Nishiuchi

## Acknowledgments

We would like to thank all members of The Genotype to Phenotype Japan (G2P-Japan) Consortium. We thank Dr. Kenzo Tokunaga (National Institute of Infectious Diseases, Japan) for sharing materials. We gratefully acknowledge the numerous laboratories worldwide that have provided sequence data and metadata to GISAID. A full list of originating and submitting laboratories for the sequences used in our analysis can be found at https://www.gisaid.org using the EPI-SET-ID: EPI_SET_230907ur.

---

**Table S1.**
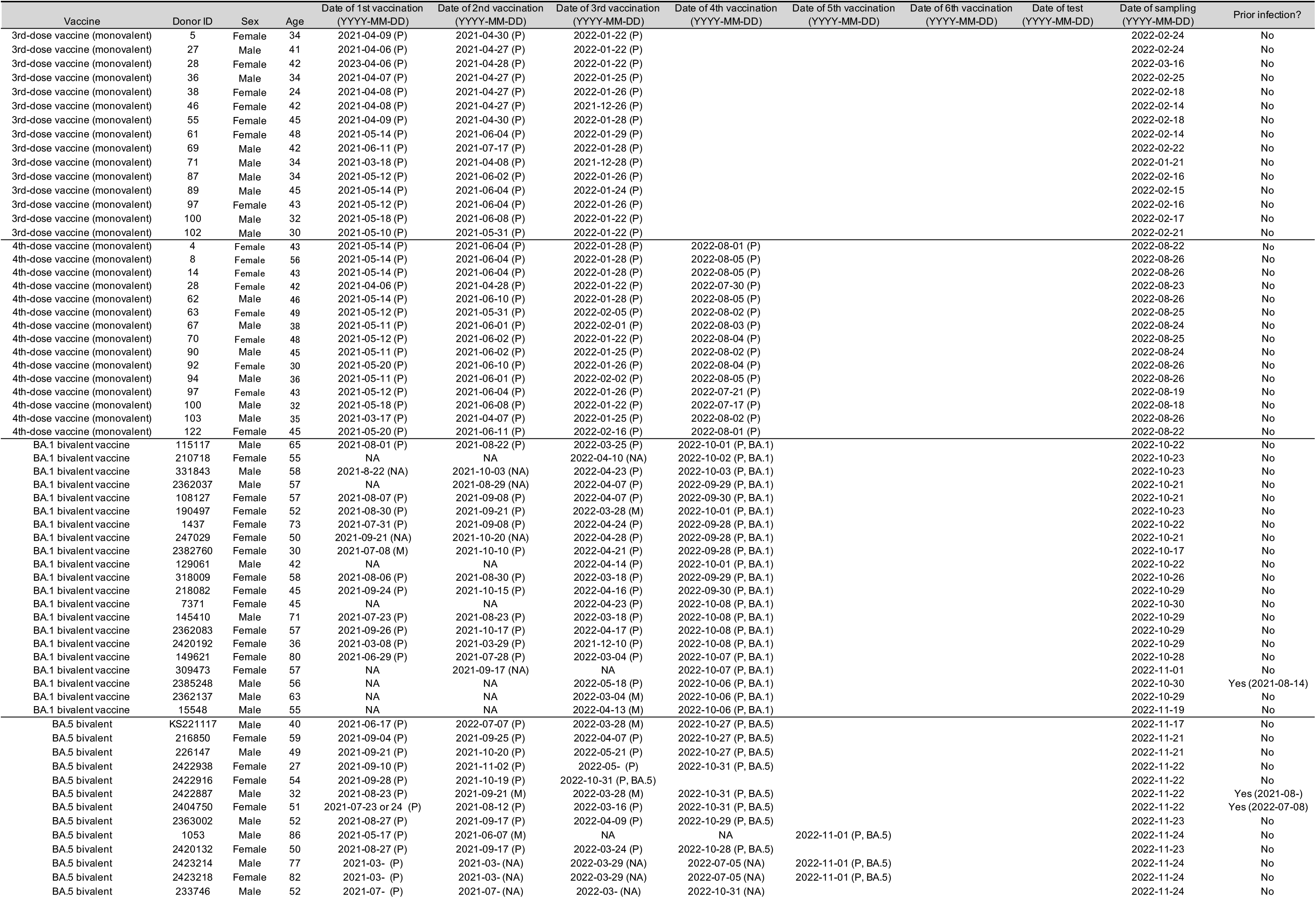
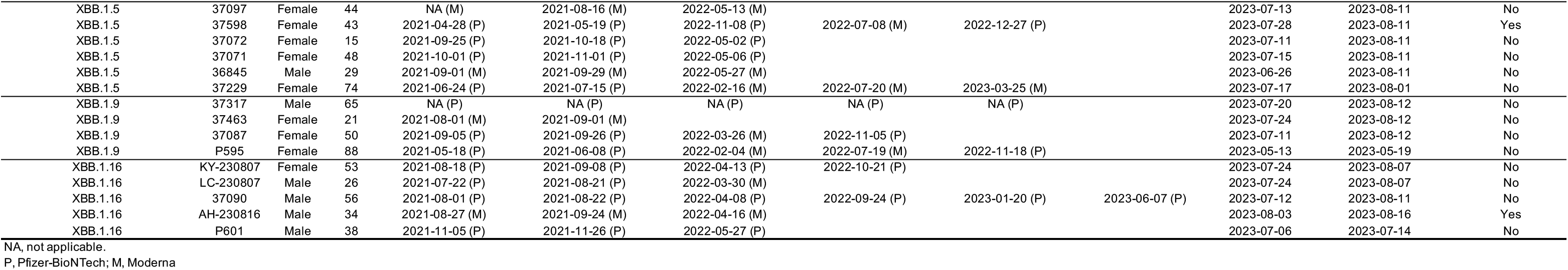
Human sera used in this study.

**Table S2.** Estimated relative Re and epidemic dynamics modeling parameters of the representative SARS-CoV-2 lineages spreading in Denmark.

| Country | PANGO lineage | Posterior mean | Posterior 2.5 percentile | Posterior 97.5 percentile | R-hat value | Effective sampling size (ESS_bulk) | Effective sampling size (ess_tail) |
| --- | --- | --- | --- | --- | --- | --- | --- |
| Denmark | BA.2.86 | 1.2882 | 1.1657 | 1.4673 | 1.0006 | 12023.7 | 8394.7 |
| Denmark | EG.5.1 | 1.1974 | 1.1732 | 1.2248 | 1.0000 | 12030.4 | 10225.5 |
| Denmark | XBB.1.16 | 1.0922 | 1.0789 | 1.1060 | 1.0006 | 11144.5 | 11542.9 |
| Denmark | XBB.1.9.2 | 1.0525 | 1.0359 | 1.0698 | 1.0002 | 12878.2 | 10880.7 |
| Denmark | XBB.1.9.1 | 1.0441 | 1.0326 | 1.0561 | 1.0003 | 11752.4 | 11594.3 |
| Denmark | EG.1 | 1.0399 | 1.0258 | 1.0540 | 1.0003 | 12465.5 | 12748.0 |
| Denmark | EU.1.1 | 1.0013 | 0.9776 | 1.0232 | 1.0001 | 14995.9 | 11025.2 |
| Denmark | FS.1 | 0.9901 | 0.9686 | 1.0106 | 1.0002 | 15092.6 | 11833.0 |
| Denmark | XBB.1.5.24 | 0.9729 | 0.9463 | 0.9977 | 1.0005 | 15121.7 | 10649.5 |
| Denmark | CH.1.1.12 | 0.9725 | 0.9425 | 0.9997 | 1.0005 | 15261.3 | 10730.2 |

The Re value of XBB.1.5 is set at 1.

**Table S3.**
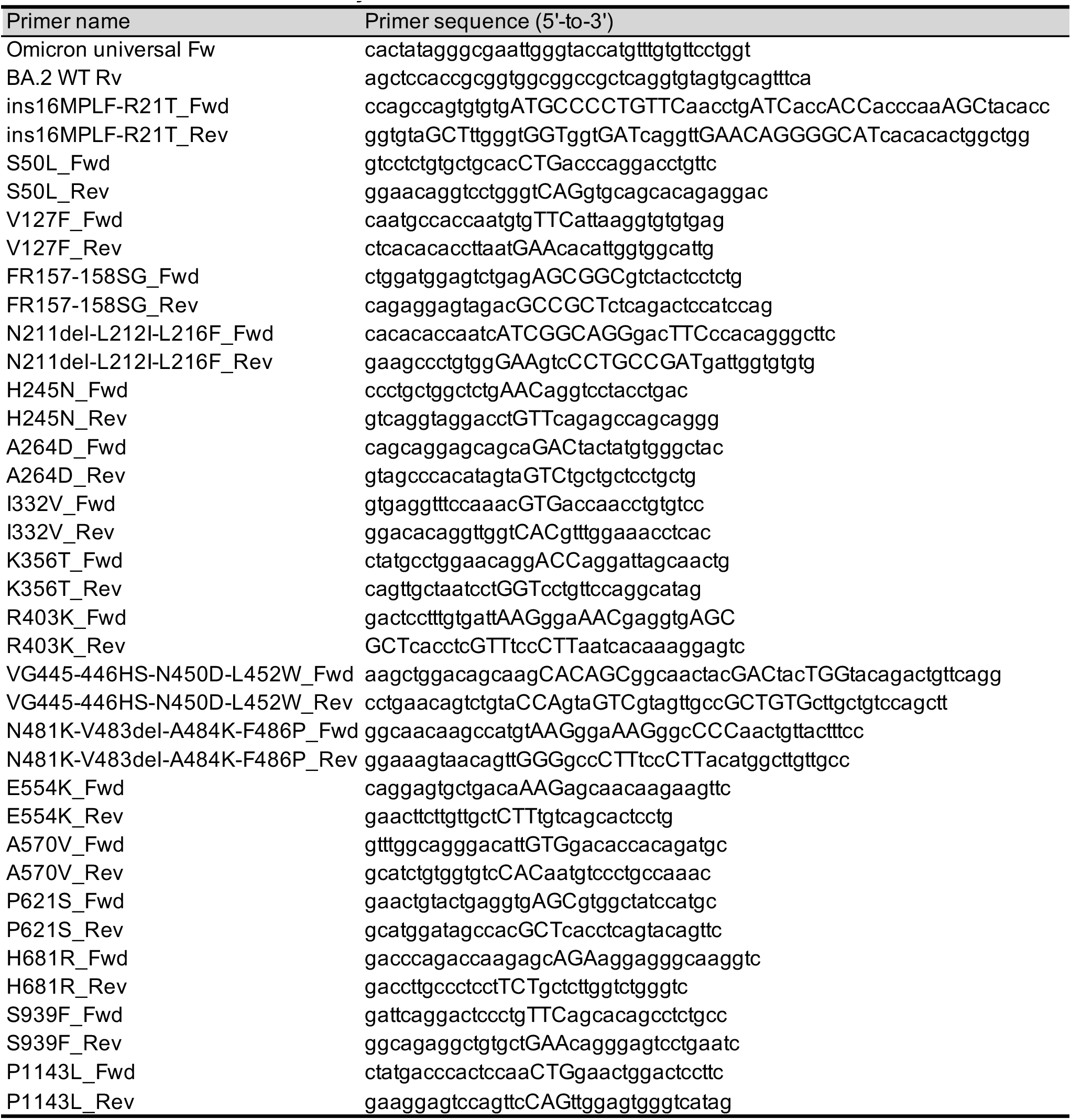
Primers used in this study.

## Notes

Conflict of interest: The authors declare that no competing interests exist.

